# An integrated computational and experimental study to elucidate *Staphylococcus aureus* metabolism

**DOI:** 10.1101/703884

**Authors:** Mohammad Mazharul Islam, Vinai C. Thomas, Matthew Van Beek, Jong-Sam Ahn, Abdulelah A. Alqarzaee, Chunyi Zhou, Paul D. Fey, Kenneth W. Bayles, Rajib Saha

**Author notes:** Corresponding author: Rajib Saha, Assistant Professor, Chemical and Biomolecular Engineering, University of Nebraska-Lincoln, Lincoln, NE-68588, USA.

## Abstract

*Staphylococcus aureus* is a metabolically versatile pathogen that colonizes nearly all organs of the human body. A detailed and comprehensive knowledge of staphylococcal metabolism is essential to understanding its pathogenesis. To this end, we have reconstructed and experimentally validated an updated and enhanced genome-scale metabolic model of *S. aureus* USA300_FPR3757. The model combined genome annotation data, reaction stoichiometry, and regulation information from biochemical databases and previous strain-specific models. Reactions in the model were checked and fixed to ensure chemical balance and thermodynamic consistency. To further refine the model, growth assessment of 1920 non-essential mutants from the Nebraska Transposon Mutant Library was performed and metabolite excretion profiles of important mutants in carbon and nitrogen metabolism were determined. The growth and no-growth inconsistencies between the model predictions and *in vivo* essentiality data were resolved using extensive manual curation based on optimization-based reconciliation algorithms. Upon intensive curation and refinements, the model contains 840 metabolic genes, 1442 metabolites, and 1566 reactions including transport and exchange reactions. To improve the accuracy and predictability of the model to environmental changes, condition-specific regulation information curated from the existing knowledgebase was incorporated. These critical additions improved the model performance significantly in capturing gene essentiality, substrate utilization, and metabolite production capabilities and increased the ability to generate model-based discoveries of therapeutic significance. Use of this highly curated model will enhance the functional utility of omics data and, therefore, serve as a resource to support future investigations of *S. aureus* and to augment staphylococcal research worldwide.

## Introduction

*S. aureus* is a versatile human pathogen that has emerged as one of the most successful infectious agents of recent times, affecting approximately 20% of the world’s population ^1-3^. The incidence of methicillin resistance at low fitness cost has significantly contributed to the rise in **c**ommunity-**a**ssociated **m**ethicillin **r**esistant ***S. aureus*** (CA-MRSA) infections, which significantly limit therapeutic options and increase rates of mortality, morbidity and costs associated with its treatment ^1,4,5^. This threat to human health has resulted in a steady interest and focus on understanding how staphylococcal metabolism relates to antibiotic resistance and pathogenesis. A number of studies have attempted to explore the metabolic aspects of antimicrobial functionality of MRSA, including nitric oxide metabolism, oxidative stress, carbon overflow metabolism, redox imbalance etc. ^6-11^. However, a complete mechanistic understanding of staphylococcal metabolism is still missing, making the identification of systematic therapeutic targets challenging.

The increase in knowledge of macromolecular structures, availability of numerous biochemical database resources, advances in high-throughput genome sequencing, and increase in computational efficiency have accelerated the use of *in silico* methods for metabolic model development and analysis, strain design, therapeutic target discovery, and drug development^12-17^. There have been a number of attempts to reconstruct the metabolism of multiple strains of *S. aureus* using semi-automated methods ^18-22^. However, the absence of organism-specific metabolic functions and the inclusion of genes without any specified metabolic functions still limit the utility of these models. These models need to be continually refined and updated to accurately predict biological phenotypes by addressing these issues as well as by reducing metabolic network gaps, elemental imbalance, and missing physiological information. Since the predictive genome-scale metabolic models of several microorganisms were useful in performing *in silico* gene essentiality and synthetic lethality analyses and yielded promising results in pinpointing metabolic bottlenecks and potential drug targets^14,23-26^, the potential for accurately modeling *S. aureus* metabolism is immense. To this end, Seif *et al* recently developed an updated genome-scale model of *S. aureus* strain JE2, incorporated 3D protein structures, evaluated gene essentiality predictions against experimental physiological data, and assessed flux distributions in different media types ^21^. Although their model was informed by multilevel omics data and a significant step toward deciphering the metabolic differences of this organism under different environmental conditions, it could further be improved by incorporating the latest annotation information, reducing the inconsistency in gene essentiality predictions, and removing spurious metabolic functionalities.

Several other studies have been dedicated to elucidating the metabolic aspects of staphylococcal virulence and to pinpoint the key metabolic “hubs” in carbon and nitrogen metabolism ^11,27-32^. However, a majority of these studies were focused on specific segments of staphylococcal metabolism and overlooked a system-wide inter-dependence that drives fitness, metabolic robustness, virulence, and antimicrobial resistance. Hence, a holistic approach of *in silico* genome-scale modeling and *in vivo* experimentation is crucial for gaining an improved mechanistic understanding of staphylococcal metabolism and, thereby, facilitating the development of novel therapeutic strategies to combat staphylococcal infections.

In this study, a comprehensive genome-scale metabolic model of *S. aureus* USA300_FPR3757 was reconstructed using annotation information from biochemical databases^33,34^ and previous strain-specific models ^19,20,34^ and validated through experimental observations and published phenotypic data. The model underwent extensive manual curation to ensure chemical and charge balance, thermodynamic consistency, and biomass precursors production. To test and inform the model, the fitness level of 1920 mutants from Nebraska Transposon Mutant Library (NTML)^35^ was assessed through an elaborate growth experiment and the metabolite excretion profiles of eight important mutants distributed across several pathways of the carbon and nitrogen metabolism were measured. The growth phenotyping results of the NTML mutants were utilized via GrowMatch procedure^36^ to reconcile *in silico* vs. *in vivo* growth inconsistencies. Upon incorporating conditional regulations in the model gleaned from existing ‘omics’ datasets^30,37,38^, the predictive capability of the model in terms of gene essentiality and metabolite excretions in different environmental conditions was further improved. Furthermore, the growth predictions from the model on 69 different carbon sources were validated against existing growth experiment^21^. Overall, this model is extensively tested by multiple available and newly-developed experimental datasets on staphylococcal metabolism and subsequently refined to pave a way forward to advance system-wide analysis of fitness and virulence.

## Results

### Reconstruction of an updated model of *S. aureus* metabolism

#### Preliminary reconstruction utilizing the existing knowledge base

A collection of 1511 metabolic reactions obtained from a consensus of recently published strain-specific models ^19,21^ was assembled into a preliminary model of *S. aureus*. Out of 842 genes in the latest strain-specific USA300_FPR3757_uid58555 model by Bosi *et al*.^19^, 109 did not have any reactions associated with them, which were not included in our model at this stage. Checking reactions from the *S. aureus* N315 model *i*SB619^20^ against the annotations of strain USA300_FPR3757 in the KEGG database^39^ resulted in the inclusion of seven unique reactions to the preliminary model. In addition, every metabolic function in the model was verified for correct gene annotations in the NCBI, KEGG, and UniProt databases and published resources^19,39-42^ to amend the model with 38 metabolic reactions and annotate 75 additional reactions with correct Gene-Protein-Reaction (GPR) rules.

These amendments resulted in a preliminary model that contained 833 metabolic genes catalyzing 1556 reactions involving 1440 metabolites. This model included reactions for central carbon metabolism, secondary biosynthesis pathway, energy and cofactor metabolism, lipid synthesis, elongation and degradation, nucleotide metabolism, amino acid biosynthesis and degradation. All the existing metabolic reconstructions of *S. aureus*^19,20,22^, including the most recently published model ^21^, used a biomass equation similar to the closely-related organisms *Bacillus subtilis*^43^ and *Escherichia coli*^44^, with additional adjustments to accommodate lipid compositions. However, *S. aureus* lacks an identifiable polyamine biosynthetic pathway and therefore cannot produce putrescine^28,45^. Therefore, putrescine was removed from the biomass equation adopted from Bosi *et al.*^19^ in the current study. Growth condition was set to glucose minimal media with other essential nutrients (see Supplementary Table 1 for details).

#### Model curation to ensure chemical balance and thermodynamic consistency

The preliminary reconstruction underwent extensive manual curation steps as outlined in the methods section. In total, 197 reactions (excluding the biomass reaction, demand, sink, and exchange reactions) were found to be imbalanced in terms of proton, carbon, nitrogen, oxygen or sulfur. Most of these reactions (*i.e*.,182 reactions) were fixed for proton imbalance and four reactions were fixed for imbalance in other elements (see Supplementary Table 2 for details). Nonetheless, a few mass- and charge-imbalanced reactions remained in the model, primarily due to the presence of macromolecules with unspecified “R”-groups and gaps in knowledge about correct reaction mechanisms. These remaining reaction imbalances are common in published genome-scale metabolic models^46^ and given that the overall stoichiometry of the reactions involving these macromolecules is correct, these imbalances do not significantly affect the performance of the model.

In addition to charge and elemental imbalances, the preliminary model had 291 reaction fluxes unnecessarily hitting the upper or lower bounds during a Flux Variability Analysis (FVA) when no nutrients were provided (see Methods section). Also, the inconsistent dissipation of ATP, which was persistent in earlier models^19,21^, also existed in the preliminary reconstruction. These two phenomena are observed when the reaction network contains thermodynamically infeasible cycles (as defined in the Methods section)^47^. To resolve these cycles, 27 reactions were made irreversible and two reactions were reversed in directionality based on available thermodynamic information and literature evidence (details in Supplementary Table 3). Furthermore, 66 reactions were turned off either due to their improper annotations or to remove lumped or duplicate reactions from the model. For example, the irreversible duplicates for several reactions including acetolactate synthase, aconitase, phosphoribosylaminoimidazole carboxylase, alcohol-NAD oxidoreductase, arginine deiminase, D-ribitol-5-phosphate NAD 2-oxidoreductase, glycerate dehydrogenase, methionine synthase, and ribokinase were removed. Also, based on available cofactor specificity information, reactions such as cytidine kinase (GTP), glycerol-3-phosphate dehydrogenase (NAD), guanylate kinase (GMP:dATP), and homoserine dehydrogenase (NADH) were turned off to ensure correct cofactor usage in these reactions. Reactions involved in polyamine synthesis and degradation were removed due to the lack of convincing evidence of polyamine functionality in *S. aureus* USA300_FPR3757 ^28,45^. After these manual curation steps, the number of unbounded reactions (reaction fluxes hitting either the upper or the lower bound without any nutrient uptake) was reduced to seven. The annotation of *S. aureus* USA300_FPR3757 genome in the KEGG database was next used to bridge several network gaps in the model. At this stage, the model contained 553 blocked reactions compared to 784 in the preliminary reconstruction. While this was a significant improvement, the model still contained a greater number of blocked reactions than other similar-sized models^21^. The blocked reactions were not removed at the current stage because they contained proper gene annotation information but either their terminal dead-end metabolite was beyond the scope of the model or no convincing evidence (*e.g*., high-score annotations) for filling the gap was available. A detailed list of the corrections and additions/removals made is given in Supplementary Table 3. The model reconstruction process, pathway distribution, and comparative model statistics are shown in Figure 1. The model is available in systems biology markup language format in Supplementary Data 1.

**Figure 1:**
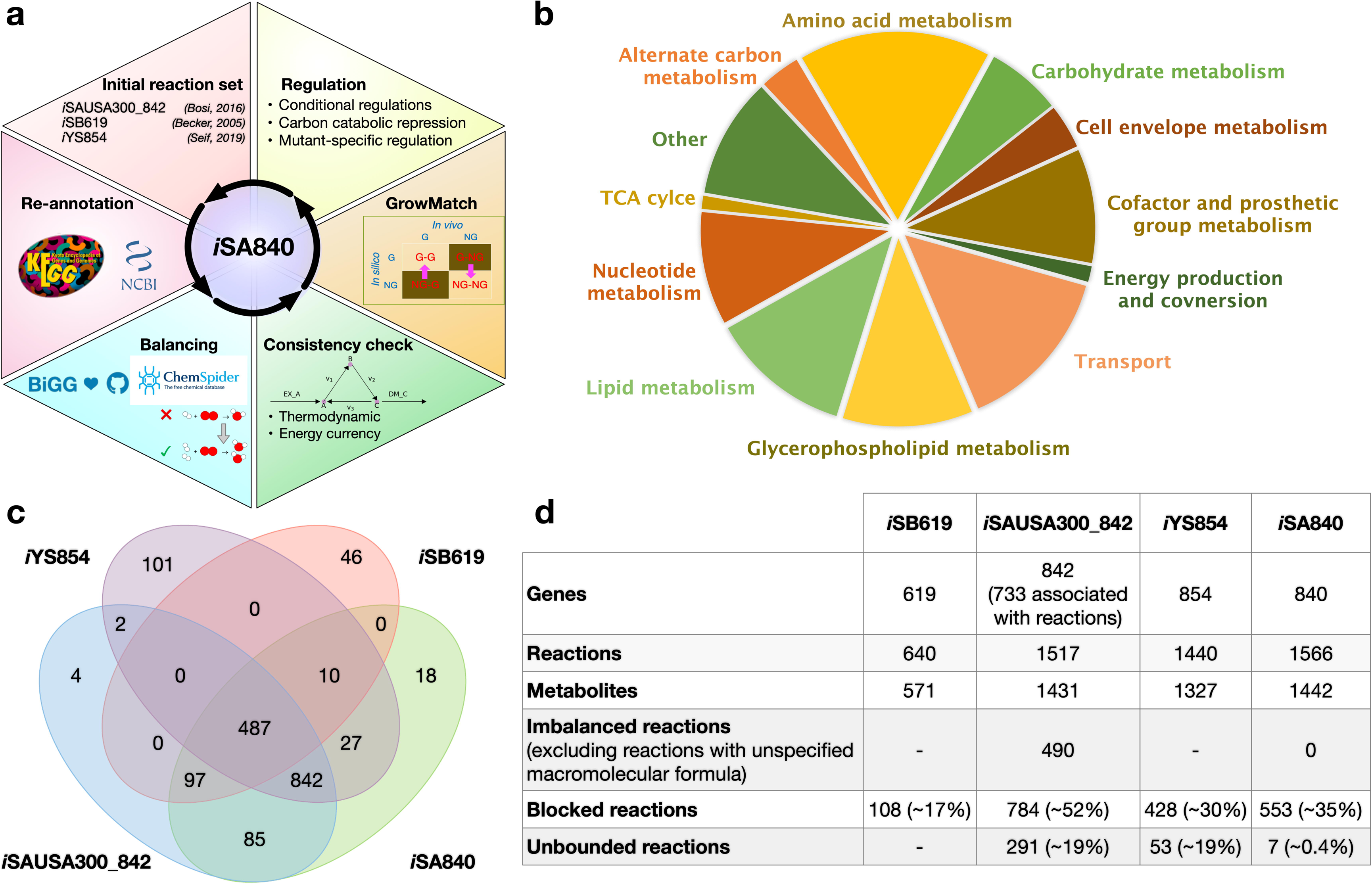
(a) The schematic of the reconstruction and curation process for *i*SA840, (b) pathway distribution of metabolic functions, (c) overlap of functionalities, and (d) comparison of model statistics with recent *S. aureus* metabolic models.

### Identifying essential genes from existing knowledgebase

We next evaluated the growth profiles of the viable *S. aureus* mutants from the NTML^35^. The variation of wild-type growth among the 384-well plates in the experiment was statistically insignificant based on z-score (see Supplementary Table 4 for detailed calculations). Out of the 1920 mutants studied, there were 154 genes whose mutations reduced growth by 10% relative to the wild-type strain and 21 mutations reduced the growth between 30% and 80% compared to the wild-type value. Out of all the genes from the NTML library, 41 genes were reported to be essential in other recent studies ^18,48-52^, whereas only 11 of them showed any significant growth inhibition (more than a standard deviation from the average wild-type growth rate) in the current study (see Supplementary Table 4 for details). Therefore, the set of essential genes was a consensus of multiple literature sources ^18,48-52^ and our current experimental study (see Methods and Supplementary Text S1 for details). Briefly, transposon mutagenesis followed by growth experiments by Valentino *et al.*^49^ and Chadhuri *et al.*^51^ identified 426 and 351 essential genes, respectively. Since the disagreement regarding gene essentiality was persistent among these datasets, the common essential gene (comprsing 319 genes) set from these two transposon mutagenesis experiment was considered to be essential, which also agreed with multiple previous growth experiments ^18,50,52^. Later, Santiago *et. al* ^48^ demonstrated that gene essentiality derived from transposon libraries can be affected by the high temperatures used to remove the plasmid delivery vehicle and also by the polar effect in disrupting expression of essential genes in the vicinity of a non-essential gene. Therefore, following their results, these false positive genes (30 in total) were excluded from the essential gene list. Finally, for the modeling purpose, only the 167 metabolic genes (excluding non-metabolic genes) present in the model were considered to be the core set of essential genes in the current study (see Supplementary Table 5 for the full list of the essential genes).

### Model refinement to reconcile growth and no-growth inconsistencies

Comparison of essential and non-essential genes between the experimental (*in vivo*) and model-based (*in silico*) gene essentiality analysis (see Methods section for details) show that there exists a significant mismatch between these two sets of results (Figure 2a). Correct model predictions for non-essential and essential genes are denoted by GG and NGNG, while wrong model predictions for non-essential and essential genes are denoted by NGG and GNG, respectively in which the first of the two terms (“G” or “NG”) corresponds to *in silico* and the second term refers to *in vivo* observations. An optimization-based procedure called Growmatch was used to reconcile the GNG inconsistencies by suppressing spurious functionalities and the NGG inconsistencies by adding miss-annotated functionalities to the model ^36^. The overall impact of applying Growmatch is shown in Figure 2b. The specificity increased from 52% to 60.5%, the sensitivity increased from 87% to 89%, and the false viability rate decreased from 48% to 39.5%. To resolve the NGG inconsistencies, metabolic functions were added from the *E. coli i*AF1260 ^44^ and *B. subtilis*^43^ metabolic models as well as the Modelseed database^34^. A total of five reactions were added to the model and three reactions were allowed to go in the reverse direction based on literature evidence or thermodynamic information (detailed procedure outlined in Supplementary Information 1), which reduced the number of NGGs by 12. Model predictions of essential genes were further improved upon the removal of spurious metabolic functions. To this end, six reactions that did not have either any gene associated with them (orphan reactions) or proper gene annotation, were removed from the model, resulting in an 18% reduction in GNGs. 81 of the GrowMatch predicted resolution strategies were not accepted because they resulted in conflicts with correct growth (GG) and no-growth (NGNG) predictions in the model. The details of the GrowMatch results are presented in Supplementary Table 5. Two example case studies for NGG and GNG inconsistency reconciliation process by GrowMatch are presented in the next section.

**Figure 2:**
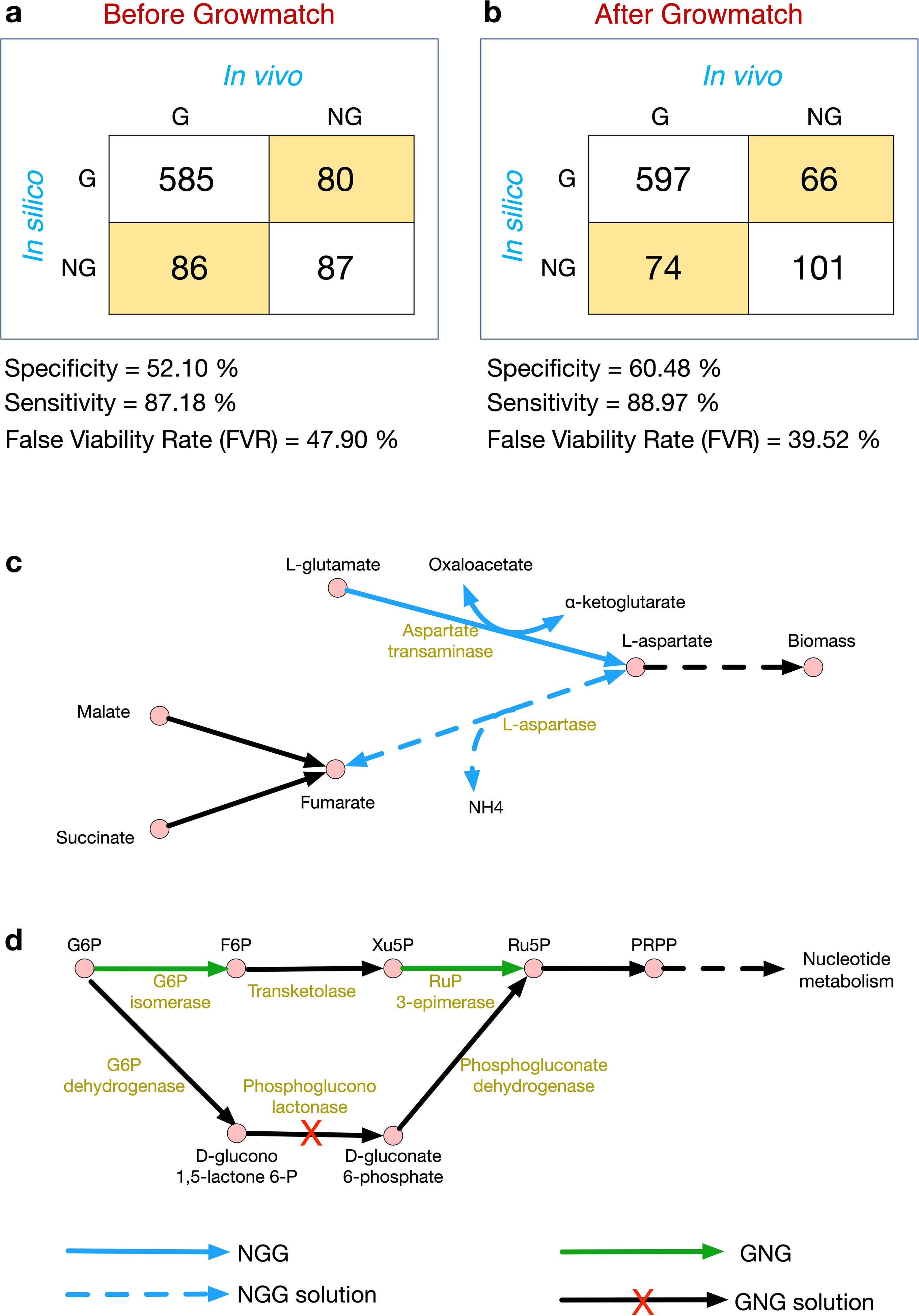
GNG table (a) before and (b) after reconciliation of growth-no growth inconsistency by GrowMatch procedure. Specificity = #NGNG/(#NGNG + #GNG), sensitivity or true viable rate (TVR) = #GG/(#GG + #NGG) and false viable rate (FVR) = #GNG/(#GNG + #NGNG), (c) a case study of NGG inconsistency and the corresponding Growmatch solution, and (d) a case study of GNG inconsistency and the corresponding Growmatch solution.

#### Case studies for reconciliation of NGG and GNG inconsistencies

The deletion of aspartate transaminase appeared to be lethal by the model prediction, whereas it was non-essential *in vivo*, making it an NGG gene (the solid blue line in Figure 2c). The addition of L-aspartase (dashed blue line in Figure 2c) rescues the growth of an aspartate transaminase deletion mutant by creating another route to generate L-aspartate, which was characterized other closely related bacteria including *E. coli* and *B. subtilis* ^53-55^. On the other hand, the Pentose Phosphate Pathway contained a GNG inconsistency, in which there were erroneous metabolic functions present in the model (Figure 2d). For example, glucose-6-phosphate isomerase and ribulose phosphate 3 epimerase are both essential genes (green highlighted genes in Figure 2d) in *S. aureus*, while they were predicted to be nonessential by the model. The reason was the presence of an alternate pathway to convert glucose-6-phosphate (G6P) to ribulose-5-phosphate (Ru5P) in the model. Since literature and database searches failed to identify the presence of phospho-glucono lactonase in *S. aureus*, it was removed, and the model was made consistent with experimental essentiality prediction of glucose-6-phosphate isomerase and ribulose phosphate-3-epimerase genes.

### Model-driven integrated study

An automated procedure like GrowMatch can significantly improve the gene essentiality predictions in the model. However, without extensive validation against experimental data and manual curations, it is difficult to obtain biologically significant and meaningful prediction capability from the model. Hence, the model was validated against multiple experimental observations from previous studies and results obtained in the current work for further refinements.

#### Incorporation of conditional regulation to enhance mutant growth predictions

The essentiality predictions for 29 amino acid catabolic pathway genes in the model was validated against the mutant growth phenotypes evaluated in a previous study^29^. The mutants were grown in a chemically defined medium (CDM) supplemented with 18 amino acids but lacking glucose. It was found that 11 of the mutations did not cause any growth defect, while 11 mutations caused intermediate growth defect and seven mutations were lethal. It was found that the model failed to recapitulate growth phenotype for nine (*ald1/ald2-* aldehyde dehydrogenase, *aspA-* aspartate aminotransferase, *gltA-* citrate synthase, *sdhA-* succinate dehydrogenase, *sdaAA/sdaAB-* serine dehydratase, *ansA-* asparaginase, *arcA1/arcA2-* arginine deiminase, and *rocF-* arginase) out of the 29 mutants, which warranted further investigation and refinements in the relevant pathways in the model. The complete growth suppression of the *pckA* mutant was not observed in the model because multiple other routes for the chemical conversion between phosphoenol pyruvate and oxaloacetate *i.e*., enolase (*eno*), phosphoshikimate 1-carboxyvinyltransferase (*aroA*) etc. are present in the model. The deletion of *ackA* gene also did not show severe growth inhibition because acetate could be generated via several routes in addition to the Pta-AckA pathway, specially *pdhABCD*, *aldA,* or *adhE.* The *gudB* mutant did not appear to be an essential gene in the model simulation because other genes including D-alanine transaminase (*dat*) and aspartate transaminase (*aspA*) could convert glutamate to alpha-ketoglutarate. However, it has been previously shown that the uptake of L-alanine in bacteria can be kinetically limited ^56^. Hence, a tighter constraint on alanine uptake was imposed in the model, which resulted in a correct prediction of the essentiality of the *gudB* gene. The essentiality of *sucC* and *sucA* genes was ensured in the model by rectifying the alternate pathway consisting of succinyldiaminopimelate transaminase (*dapE*) and tetrahydrodipicolinate succinylase (*dapD*). In addition to that, the TCA cycle reactions converting citrate to succinyl-CoA were constrained to allow flux towards the forward direction only. Two of the gaps in the histidine transport pathway and proline catabolism were filled during the refinement process to allow for utilization of these alternate carbon sources in the absence of glucose. Ornithine-putrescine antiport, lactate dehydrogenase (ferricytochrome), malic enzyme (NADP), succinyldiaminopimelate transaminase etc. were removed from the model due to the lack of evidence of these functionalities in *S. aureus*. Upon these refinements, the model was able to correctly predict 24 (out of 29) of the mutant phenotypes. The model refinements in the central metabolic pathway in terms of correction of reaction directionality, additions, and deletions are shown in Figure 3.

**Figure 3:**
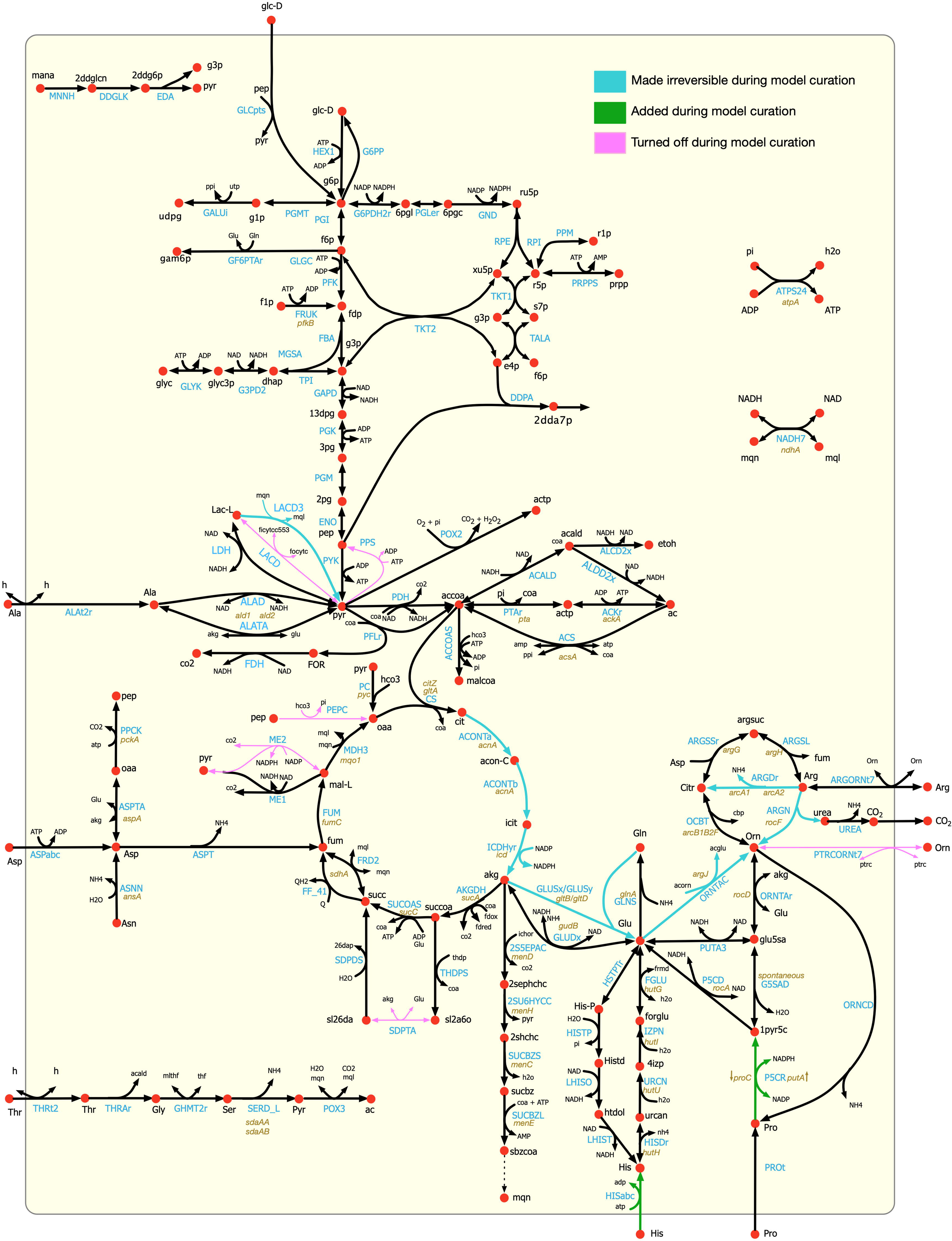
Refinements in the central metabolic pathway of the model *i*SA840 showing correction of reaction directionality, additions, and deletions.

#### Metabolite excretion profiles of mutants with altered carbon metabolism

In addition to the model refinements mentioned in the preceding section, we determined the metabolite excretion profiles of eight mutants during exponential growth (Figure 4 and Supplementary Table 6) and compared them to the model predicted excretion patterns in both CDM and CDMG (CDM media with added glucose) media. The mutants considered were *pyc* (pyruvate carboxylase), *citZ* (citrate synthase), *sucA* (2-oxoglutarate dehydrogenase), *ackA* (acetate kinase), *gudB* (glutamate dehydrogenase), *ndhA* (NADH dehydrogenase), *menD* (menaquinone biosynthesis protein) and *atpA* (a subunit of ATPase). These mutants were selected for their potential in identifying carbon and nitrogen redirection pathways as they affect important metabolic pathways associated with central metabolism including glycolysis, TCA cycle, gluconeogenesis, Electron Transport Chain (ETC), cellular redox potential and overflow metabolism. In general, supplementation of glucose (CDMG) as the primary carbon source resulted in the excretion of acetate as the major byproduct in all mutants (Figure 4). In CDM, the *ackA*, *gudB, ndhA, atpA*, *and menD* mutants displayed delayed growth kinetics (data not shown). Although acetate remained a major byproduct of strains in CDM, this was due to amino acid deamination as evidenced by ammonia excretion (Figure 4). As carbon flux through the ATP-generating Pta-AckA pathway is significant in *S. aureus*, we also observed the excretion of pyruvate and redirection of carbon flux towards acetoin and α-ketoglutarate in the *ackA* mutant (Figure 4). Mutations that affected respiration (*ndhA* and *menD*) of *S. aureus* resulted in increased levels of lactate production to maintain cellular redox when grown in CDMG (Figure 4). The disruption of ATP production due to mutation of *atpA* was offset by increased acetate production and glucose consumption. The increased flux of glucose through the Pta-AckA pathway to generate acetate likely compensated for the decrease in ATP production due to a faulty ATPase.

**Figure 4:**
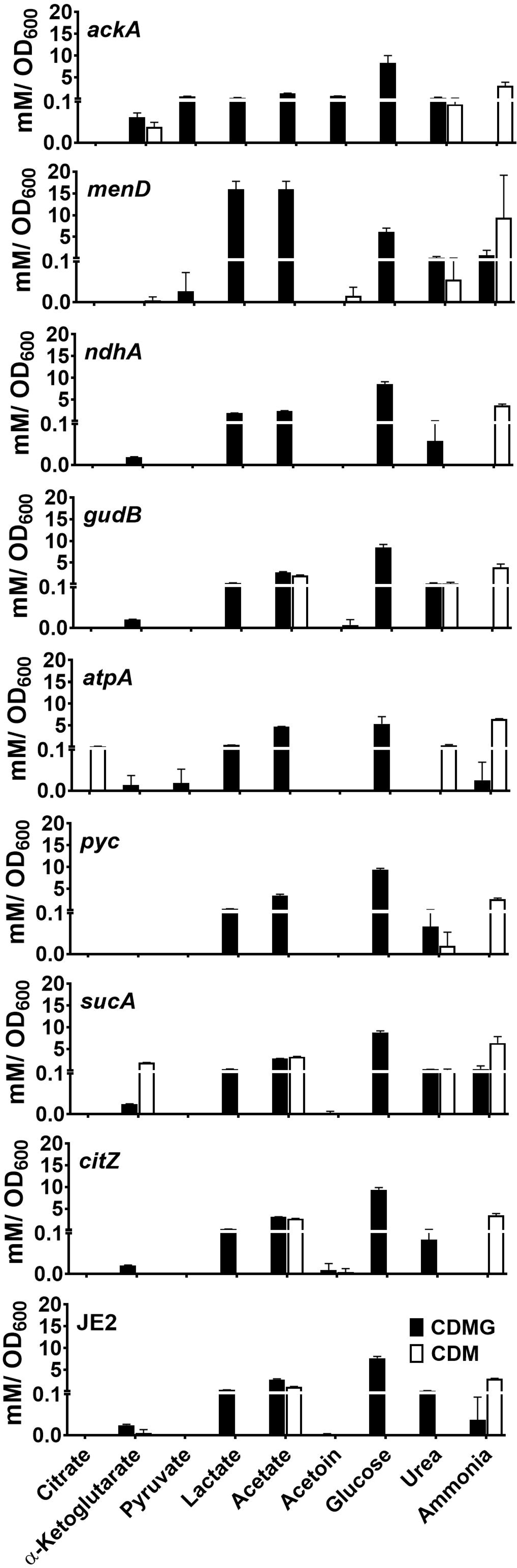
Metabolite excretion profile of multiple *S. aureus* mutants with altered carbon and nitrogen metabolism.

The comparison of experimental results and model predictions revealed multiple inconsistencies that motivated an extensive search for additional metabolic regulations in *S. aureus* in different media types. The full list of regulations can be found in Supplementary Table 7. A major regulatory system that was incorporated in the model was the carbon catabolite repression, which is a well-studied global regulatory process in low-GC Gram-positive bacteria in the presence of a preferred carbon source that induces the repression of genes involved in the metabolism of alternative carbon sources ^30^. CcpA, the carbon catabolite control protein, is known to repress genes involved in the utilization of amino acids as alternative carbon sources in the presence of glucose^38^. In addition, SrrAB and Rex-dependent transcriptional regulation are prominent driving forces of metabolic flux through respiratory metabolism that was integrated into the model. Furthermore, mutant-specific repression of respiration, histidine and ornithine metabolism, and pyruvate metabolism were imposed on the model for the *menD* mutant. For each of the mutants, the incorporation of these regulations resulted in a deviation of the metabolic flux space (defined as the range between the minimum and maximum flux through reactions, see Methods section for details) compared to the wild type, as illustrated in Figure 5.

**Figure 5:**
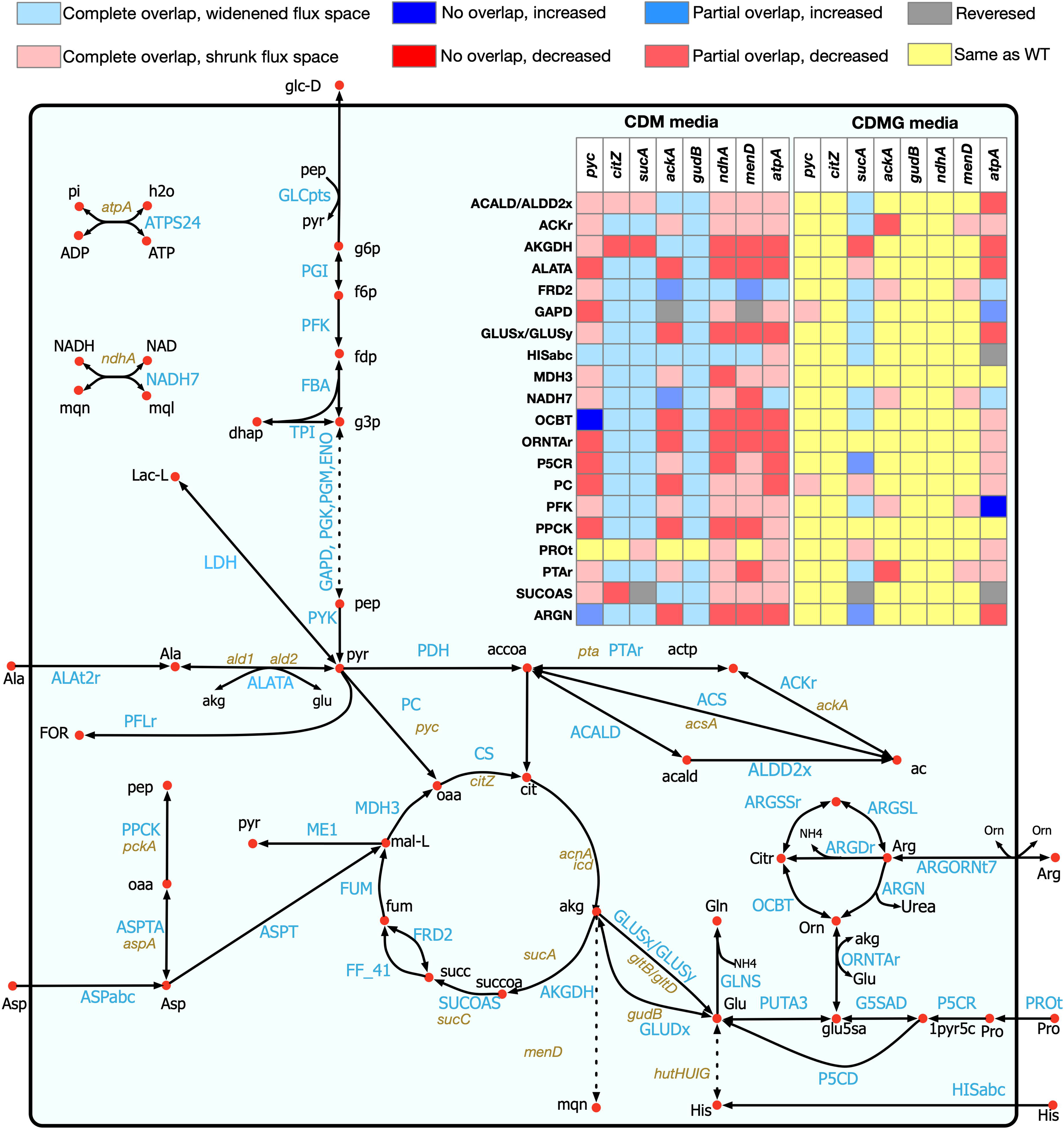
Shifts in flux space for eight mutants in the central carbon and nitrogen metabolic pathway. Every row in the table (inset) denotes a reaction as identified in the pathway map. The relative shifts compared to the wild type flux space are color-coded according to the legend in the figure.

Among the eight mutants, the model-predicted excretion patterns for acetate and lactate in *sucA* and *ackA* mutants agreed with the experimental results of decreased excretion in CDMG media, compared to the wild type strain. In CDM media, while no significant change in lactate excretion was observed, acetate excretion was decreased in the *ackA* mutant compared to the wild-type strain, due to inactivation of the Pta-AckA pathway. On the other hand, the *sucA* mutant in CDM media showed increased production of acetate due to increased flux space in the Pta-AckA pathway (see Figure 5). The Pta-AckA pathway is known to supply a major portion of the ATP required for growth ^27^. With the *atpA* gene turned off in the model Pta-AckA pathway supplied most of the ATP demand, which increased the acetate production in CDMG media for the *atpA* mutant compared to the wild-type. However, in CDM media, the model could not sustain the ATP maintenance demand of the *atpA* mutant and therefore, did not produce any acetate. In CDMG media, the model-predicted excretion profile for urea in all of the mutants matched with the experimental observations. In CDM media, the model predictions of higher urea excretion compared to the wild-type strain agreed with the experimental observations for *pyc*, *gudB*, *ndhA*, and *menD* mutants. Similar to the experimental results, excretion of ammonia was predicted by the model in all mutants when glucose was absent (CDM media). These correct predictions can be attributed to the deamination of the amino acids consumed in CDM media when the cell adapts to amino acids due to CcpA-mediated control of amino acid metabolism.

The incorporation of regulatory information improved the predictive capabilities of other mutants. For example, incorporation of regulation based on the Rex and SrrAB repressors’ effect on central carbon metabolism allowed the model to correctly simulate the oxygen deprivation in the model, which, in turn, resulted in correct predictions of decreased acetate excretion by the *ndhA* mutant in both CDM and CDMG media. Rex and SrrAB-mediated repression of pyruvate formate lyase (PFLr), alcohol dehydrogenase (ACALD, ALDD2x) and other pathways downstream of pyruvate shifted carbon flux away from the acetate production. At the same time, the flux space for lactate dehydrogenase (LDH) widened, which allowed for more lactate excretion in the CDMG media. In the *menD* mutant, mutant-specific regulation information from Kohler *et al* ^37^ resulted in the correct prediction of lactate and acetate excretion. A mutation in *menD* or any other gene in the menaquinone biosynthesis pathway resulted in weakened respiratory functions and emulated anaerobic condition in the cell, which in turn caused a significant increase in the excretion of lactate in CDMG media. However, although the respiratory functions were downregulated in CDM media (apparent from the shrinkage of the flux space), there was no change in acetate excretion compared to the wild type strain. In CDMG media, the conversion of pyruvate to oxaloacetate by pyruvate carboxylase was not active in the wild type model. A small amount of phosphoenol pyruvate was converted to oxaloacetate (via PEPC), which was then used in the conversion of glutamate to aspartate. However, since no convincing evidence for phosphoenol pyruvate carboxylase was found in *S. aureus*, PEPC was removed. This refinement shifted carbon flux through pyruvate carboxylase in the wild type model and also resulted in correct model prediction of acetate excretion in CDMG media when pyruvate carboxylase was turned off.

While the incorporation of the CcpA, Rex and SrrAB regulations was critical in capturing the physiological behavior of *S. aureus* by the model, it should be noted that there are still gaps in our knowledge about the quantitative repression effect on the reaction fluxes in the presence of these regulators. For example, in CDMG media, ammonia production was not predicted in the *menD*, *atpA,* and *sucA* mutants by the model, which was observed experimentally. However, upon further investigation, it was observed that relaxing the repressions of reaction fluxs that were imposed on the model due to CcpA, Rex, and SrrAB regulators, the discrepancies were removed. In CDMG media, the *citZ* mutant correctly predicted the excretion pattern of acetate, because with the reduced flux space for the TCA cycle reactions, more carbon could be directed to the Pta-AckA pathway. However, in the CDM media, when amino acids were the primary source of carbon, deletion of the *citZ* gene did not have any effect on the model predicted flux space in the Pta-AckA pathway. In the *pyc* mutant, carbon flux to oxaloacetate was directed through malate dehydrogenase (MDH3) in the model, which involved the consumption of menaquinone produced by cytochrome oxidase BD. When the *pyc* gene was active, the same conversion was mediated through malic enzyme (ME1) and the pyruvate carboxylase (PC). However, since the model could accommodate the metabolic shift in both the wild-type and *pyc* mutant, no change in the excretion rate of acetate or lactate was observed. Also, while the CcpA repression was active, the deletion of the *gudB* gene in the model did not have any effect on the lactate and acetate excretion profiles in CDMG media. In CDM media, the model prediction for no lactate production was consistent with experimental observations but still no effect was observed on acetate production. Also, the model predicted a lower urea production rate in the *atpA* mutant compared to the wild-type strain, while it was higher in our experiments. Also, no urea excretion was observed in the *citZ* mutant in our experiments but model predicted urea excretion at the same rate as the wild-type strain. The reason for these inconsistencies could be the lack of a complete understanding of the regulatory processes that affects the relationship between amino acid catabolism, urea cycle, TCA cycle and pyruvate metabolism. These inconsistencies warrant further investigation into CcpA-mediated metabolic control.

#### Estimation of carbon catabolism capacity of the model

In order to further test the accuracy of the model, the growth predictive capability of the model was validated against a recent study of carbon source utilization by *S. aureus* strain USA300-TCH1516 by Seif *et al*.^21^. Out of the 69 carbon sources tested, the authors observed growth on 53 metabolites and no growth on 16 metabolites in their BIOLOG experiment. Our model correctly predicted growth on 41 and no-growth on 12 of the carbon sources, and falsely predicted growth on four and no-growth on 12 carbon sources (see Supplementary Table 8 for details). In comparison, *i*YS854 correctly predicted growth on 42 and no-growth on 5 of the carbon sources, and falsely predicted growth on 11 and no-growth on 11 carbon sources. Overall, our model achieved a specificity of 75%, a precision of 91%, and an accuracy of 77%, which in general are either at par with or better than previously developed models^21^ and further demonstrates the improved predictive capability of this new model.

## Discussions

In the current study, an updated and comprehensive genome-scale metabolic model of the methicillin-resistant human pathogen *S. aureus* USA300_FPR3757 was reconstructed from the previous strain specific models ^19-21^, amended using annotations based on KEGG database^39^, and refined based on published and new experimental results. Reactions were examined and fixed to ensure chemical and charge balance and thermodynamic consistencies. The extensive manual curation performed on the preliminary reconstruction resulted in improved prediction capabilities and successful capture of experimentally observed metabolic traits. All these demonstrate the necessity of exhaustive manual scrutiny and rectification of automated reconstructions. The growth and no-growth analysis and the resolution of inconsistencies between *in silico* growth predictions and *in vivo* results using the Growmatch algorithm ^36,57^ reinforces the importance of the iterative procedure of model refinement using experimental observations. Further experimental results from mutant growth and metabolite excretion studies enabled high-resolution model refinements to further enhance the predictive capabilities of the model. The final genome-scale metabolic reconstruction (*i*SA840) is therefore a product of the series of automated and manual curation steps.

Our growth evaluation experiment revealed varying degrees of growth inhibition of the NTML mutants compared to the wild type strain and identified subtle disagreements in gene essentiality predictions of other studies ^18,48-52^. Therefore, the true set of essential genes required further scrutiny, which is why, as a conservative estimate, we used a consensus set of essential genes by utilizing the existing knowledge base and our own experimental findings (more details in Supplementary Information 1). Moreover, several mutants compromised in growth could be found in all the different methods, which did not appear to inhibit growth significantly during model simulations. Instead, the model either predicted growth at full capacity or became completely growth-inhibited. This phenomenon suggests that the model has degeneracy in the flux space that may compensate for lost functionality by redirecting or shifting metabolic fluxes. This issue calls for a more rigorous study of the regulatory influences and necessitates further future studies in enzymatic efficiencies and kinetics associated with important metabolic pathways.

The growth phenotyping studies of mutations in the amino acid catabolic pathway^29^ revealed shifts in *S. aureus* metabolism in the absence of a preferred carbon source and elucidated the extent of carbon catabolic repression, which allowed us to make necessary amendments to the model in terms of correction of reaction directionality, removal and addition of reactions, and specifying cofactor utilization across the central metabolic pathway (see Figure 3 for details). The change in media components (CDM vs. CDMG) resulted in a significant redistribution of metabolic flux in the model, as was evident from the shifts in flux space for different mutants in the carbon and nitrogen metabolic pathways. These shifts predicted how inactivation and/or repression of TCA cycle, respiration, electron transport and ATP generation could impact the cellular redox balance, metabolite production, and fitness. While the model predictions for acetate and lactate production in the *ackA* and *sucA* mutants and ammonia and urea production in *ackA*, *pyc*, *gudB*, *ndhA*, and *menD* mutants matched with experimental results, other mutants showed deviations in their metabolite excretion behavior. The prediction capability of the model was improved upon the addition of regulatory information obtained from existing ‘omics’ datasets^30,37,38^. For example, incorporation of Rex and SrrAB regulation caused repression on pyruvate metabolism and alcohol dehydrogenase pathways, which resulted in correct predictions of acetate excretion by the *ndhA* mutant in both CDM and CDMG media, and by the *citZ* and *pyc* mutants in CDMG media. Moreover, imposing mutant-specific repressions was critical to achieving predictive results for the acetate and lactate excretion in the *menD* mutant and ammonia and urea excretion in the *atpA* mutant. However, the current knowledge of the regulatory landscape in *S. aureus* is not sufficient to explain some of the inconsistent metabolite production trends in the remainder of the mutants, thus, warranting the need for further investigation.

*S. aureus* remains a significant threat to human health, which drives a growing number of studies towards understanding how staphylococcal metabolism relates to antibiotic resistance and pathogenesis. Very few studies have addressed these interrelationships from a systems biology perspective, which requires a predictive *in silico* metabolic model capable of capturing the biochemical features of the pathogen. This work addresses these gaps through the development of a detailed metabolic model informed not only from existing resources, such as the NTML, *in silico* genome sequences, annotation databases, and theoretical metabolic stoichiometry but also from our own experimental studies on mutant fitness, gene essentiality, and metabolite excretion profile. The results presented in this work demonstrate the predictive capacity of the new genome-scale metabolic reconstruction of *S. aureus* USA300_FPR3757, *i*SA840, in different environments, utilizing different substrates, and with perturbed genetic contents, which paves the way for a mechanistic understanding of *S. aureus* metabolism. This latest genome-scale model of *S. aureus* demonstrates high performance in capturing gene essentiality, mutant phenotype and substrate utilization behavior observed in experiments. However, the accuracy and prediction capability, as well as the ability to generate model-based drug-target discoveries, can be further enhanced by incorporating extensively vetted flux measurements, quantitative proteomics, and kinetic measurements of metabolic intermediates. The development of a more accurate systems-level metabolic model for *S. aureus* will have a tremendous impact on future scientific discoveries and will be a valuable resource shared among the staphylococcal research community for the identification and implementation of intervention strategies that are successful against a wide range of pathogenic strains.

## Methods

### Preliminary model reconstruction and curation

#### Preliminary model and flux balance analysis

The primary reaction set was obtained from the genome-scale metabolic reconstruction of *S. aureus* USA300_FPR3757 by Bosi *et al.* ^19^ and a recent model of strain JE2 by Seif *et al.* ^21^. Reactions from the *S. aureus* N315 model *i*SB619 ^20^ were checked against annotations of *S. aureus* USA300_FPR3757 based on the KEGG database^39^ and merged with the reaction set to get the preliminary model. Flux balance analysis (FBA)^58-60^ was employed during model testing, validation, and analyzing flux distributions at different stages of the study. For performing FBA, the reconstruction was represented in a mathematical form of stoichiometric coefficients (known as stoichiometric matrix or S-matrix), where each column represents a metabolite and each row signifies a particular reaction. In addition to the mass balance constraints ^61^, environmental constraints based on nutrient availability, the relational constraint of reaction rates with concentrations of metabolites, and thermodynamic constraints were imposed as necessary. The effects of gene expressions were incorporated as regulatory constraints on the model as the cell adapted to change in media or gene knockouts ^62^. The non-growth-associated ATP maintenance demand was estimated to be 5.00 mmol/gDCW.hr in CDM media and 7.91 mmol/gDCW.hr in CDMG media in this study, according to the established protocol^63^.

#### Rectification of reaction imbalances

To ensure that each of the reactions in the model is chemically balanced, the metabolite formula and the stoichiometry of the reactions were checked against biochemical databases ^34,39,64,65^. For balancing the reactions imbalanced in protons, the protonation state consistent with the reaction set in the preliminary model was checked and additions/deletions of one or multiple protons or water on either the reactant or the product side were performed. For the other elements, correct stoichiometry was incorporated into the S-matrix. Reaction with unspecified macromolecule formula were not rectified.

#### Identification and elimination of thermodynamically Infeasible Cycles

One of the limitations of constraint-based genome-scale models is that the mass balance constraints only describe the net accumulation or consumption of metabolites, without restricting the individual reaction fluxes. Therefore, they have an inherent tendency to ignore the loop low for electric circuits which states that there can be no flow through a closed loop in any network at steady state ^47^. While biochemical conversion cycles like TCA cycle or urea cycle are ubiquitous in a metabolic network model, there can be cycles which do not have any net consumption or production of any metabolite. Therefore, the overall thermodynamic driving force of these cycles are zero, implying that no net flux can flow around these cycles ^47^. It is important to identify and eliminate these Thermodynamically Infeasible Cycles (TICs) to achieve sensible and realistic metabolic flux distributions.

To identify Thermodynamically Infeasible Cycles in the model, all the nutrient uptakes to the cell were turned off and an optimization formulation called Flux Variability Analysis (FVA) was used^66^. FVA maximizes and minimizes each of the reaction fluxes subject to mass balance, environmental, and any artificial (*i.e*., biomass threshold) constraints ^66^. The reaction fluxes which hit either the lower bounds or upper bounds are defined as unbounded reactions and were grouped as a linear combination of the null basis of their stoichiometric matrix. These groups are indicative of possible thermodynamically infeasible cycles. To eliminate/destroy the cycles, duplicate reactions were removed, lumped reaction were turned off or reactions were selectively turned on/off based on available cofactor specificity information (see Supplementary Information 1 for details).

### Evaluation of growth profiles of mutants in NTML

Pre-cultures of wild-type and isogenic transposon mutant strains were grown overnight aerobically in 384-well plates containing 100 μL of Tryptic Soy Broth (TSB)/ well. The overnight cultures (1 μL) were seeded into a fresh 384-well plate containing TSB (100 μL/ well) using a solid 384 pin tool (V & P Scientific) and cultured for 24 h at 37°C under maximum agitation in a TECAN microplate reader. Growth was monitored by recording the optical density (OD_600_) of cultures for 24 h at 30-minute intervals. The area under the growth curve (AUC) was calculated as a measure of growth for each strain and used for comparative analyses.

### Elimination of Growth and No-growth Inconsistencies between model predictions and experimental data

#### Gene essentiality analyses

Metabolic robustness of an organism in the event of genetic manipulations are attributed to the essentiality of the respective gene(s) under a specific nutrient medium or regulatory condition ^24^. In any metabolic reconstruction, there are either missing necessary functionalities in the model or erroneous pathways present in the model, mainly due to missing or wrong annotation information. To identify these inconsistencies in the model, *in silico* essential and non-essential genes were identified by turning off the reaction(s) catalyzed by the gene following the Boolean logic of the Gene-Protein-Reaction (GPR) relationships, and estimating growth as a result of the deletion. Isozymes (i.e., proteins/genes with an “OR” relationship) for essential reactions are not considered as essential, and for reaction catalyzed by protein with multiple subunits (i.e., proteins/genes with an “AND” relationship), each gene responsible for each subunit is considered essential. A mutant was classified as lethal if its growth rate is below the threshold of 10% of the wild type growth rate.

*In vivo* essential genes were curated from multiple sources ^18,48-52^, as explained in detail in the Supplementary Text S1. Most of the essential genes were determined by randomly inserting transposons into *S. aureus* and excluding mutations that remained after growing the cells ^48,49,51^. An adaptation of data from multiple sources using antisense RNA was also used to determine essential enzymes and thus essential genes through the Boolean relationships ^18,50,52^. Genes reported to be essential in any sources were considered essential unless there was evidence suggesting otherwise ^18,48-52^. There were three types of positive evidence. First, mutants obtained from Nebraska’s Transposon Mutant Library ^35,67^ were not considered essential unless it was found to be domain-essential ^48^. This is because the transposon may have inserted in a non-essential part of the gene, allowing a partially functional protein to be formed. Second, if the gene was found to be essential at only 43°C, then it is evident that the gene was incorrectly found to be essential in literature because of a high-temperature plasmid curing step in the processes used in the other literature sources ^48^. Third, if the gene was found to be essential using a promoterless transposon insert, but not with promoter-containing methodologies, then the gene is upstream of an essential gene, and other sources found it to be essential due to polar effects that disrupt expression ^48^. The step-by-step methodology used in determining core essential gene set is illustrated in Supplementary Figure S4.

Out of the concensus set of the essential genes, 167 metabolic genes that are present in the *i*SA840 metabolic model were considered for further model refinements. The results of the *in silico* growth estimation were compared with these experimental evidences, and the genes were classified based on the matches and mismatches between *in silico* and *in vivo* results. Correct model predictions for non-essential and essential genes are denoted by GG and NGNG, while wrong model predictions for non-essential and essential genes are denoted by NGG and GNG, respectively. GNG inconsistencies imply that the metabolic model erroneously contains reactions that complement for the lost gene function. In contrast, NGG inconsistencies are generally indicative of missing or poor annotations in the model.

#### Using GrowMatch to resolve inconsistencies

To resolve the growth and no-growth inconsistencies in the model, an automated procedure called GrowMatch was used^36^. GrowMatch tries to reconcile GNG predictions by suppressing spurious functionalities that were mistakenly included in the model and NGG predictions by adding missing functionalities to the model while maintaining the already identified correct growth and no-growth predictions ^36^. Every suggested GrowMatch modification was filtered for the resolution of conflict following the procedure of Henry *et al*. in 2009 ^43^. A detailed explanation of these cases can be found in the Supplementary Table 5.

### Determination of metabolite excretion profiles of mutants

To determine the metabolite excretion profile of various strains, cell-free culture supernatants were analyzed by HPLC for multiple weak acids, acetoin, and sugars as previously described. Briefly, the analysis was performed isocratically at 0.5 mL/min and 65°C using a Biorad Aminex HPX-87H cation exchange column with 0.13N H_2_SO_4_ as the mobile phase. The peaks corresponding to various metabolites were identified by their retention time obtained by using genuine standards. Absolute concentrations were determined from calibration curves specific to each metabolite. Ammonia and urea were measured using a kit (R-biopharm) according to the manufacturer’s protocol.

### Incorporation of regulation in the model

Regulation information for *S. aureus* in terms of differential expression of genes or high/low abundance of the corresponding proteins were accumulated from multiple sources as listed in Supplementary Table 7. Gene-Protein-Reaction (GPR) Boolean relationships for each of the genes were used to determine the corresponding reactions to be regulated in model simulations in different conditions. If a reaction in catalyzed by multiple isozymes, the reaction was only suppressed if all of the isozymes were downregulated in a certain condition. For a reaction catalyzed by multiple subunit proteins, it was suppressed if any of the genes responsible for a subunit was downregulated. For aerobic vs. anaerobic simulations in the model, the lower bound and upper bound for the regulated reactions were set to zero. For CcpA, SrrAB, and Rex repression, the allowable flux ranges were limited to 50% of their wild-type flux values. For the reactions suppressed in *menD* mutant, a similar flux limitation was imposed.

## Data Availability

All data generated or analyzed during this study are included in this published article and its supplementary information files.

## Competing Interests

The authors declare no competing interest for the presented work.

## Contributions

R.S., V.C.T., P.D.F. and K.W.B. conceived the study. R.S. and V.C.T supervised the study. V.C.T., J.S.A, A.A.A and C.Z. performed and analyzed the *in vivo* studies. M.M.I performed the *in silico* experiments and analyses. M.V.B. and M.M.I. developed the required software programs, models, and graphics. M.M.I., M.V.B., R.S., and V.C.T. wrote the manuscript. All authors have reviewed and approved the submission of the manuscript.

## Funding

This work was supported by the Nebraska Systems Science Initiative Seed Grant [**WBS# 21-3209-0013**] to R.S., V.C.T, K.W.B., and P.D.F.; NIH/NIAID P01AI083211 to K.W.B. and P.D.F; NIH/NIAID R01AI125588 to V.C.T.

## Supplementary Information

**Supplementary Information 1:** Strategies for fixing thermodynamically infeasible cycles, consensus of *in vivo* gene essentiality information, and details of GrowMatch procedure and results.

**Supplementary Table 1**: Growth medium definition.

**Supplementary Table 2**: Fixed reactions imbalanced in carbon, hydrogen and oxygen.

**Supplementary Table 3**: All the reactions turned off or directionality changed or removed as a duplicate during model curation steps.

**Supplementary Table 4**: NTML mutant growth data and statistical analysis.

**Supplementary Table 5**: Gene essentiality information, comparison of model and experimental essentiality results, GNG tables, Growmatch results, and rejected Growmatch suggestions.

**Supplementary Table 6**: Metablite excretion profiles by mutants in CDM and CDMG media.

**Supplementary Table 7**: Regulations and repressions imposed on the model.

**Supplementary Table 8**: Model predictions on utilization of different carbon sources and comparison with BIOLOG experimental results.

**Supplementary Data 1**: The *S. aureus* USA300_FPR3757 metabolic model in systems biology markup language format.

Supplementary information is available at NPJ Systems Biology and Applications’ website.

