## Supplementary Information 1 for "An integrated computational and experimental study to elucidate *Staphylococcus aureus* metabolism"

**Supplementary material**

1. **Strategies for fixing thermodynamically infeasible cycles**

To fix the thermodynamically infeasible cycles in the models, three distinct cases were addressed.

**Case 1: Duplicate reactions that run in opposite direction**

In this case the model contains duplicates of the same reaction, often one being irreversible and one being reversible. The cycle can be broken by removing or turning off one of the reactions, usually the irreversible one if no concrete thermodynamic information is available.

Example:

Phosphoglycerate dehydrogenase (PGCD): nad_c[c] + 3pg_c[c] -> h_c[c] + nadh_c[c] + 3php_c[c]

and

Phosphoglycerate dehydrogenase reversible (PGCDr): nad_c[c] + 3pg_c[c] <=> h_c[c] + nadh_c[c] + 3php_c[c]

Solution:

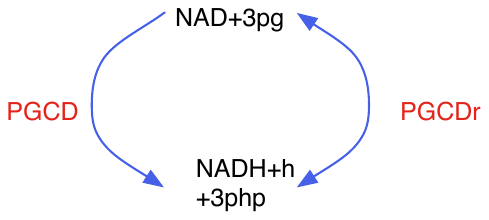

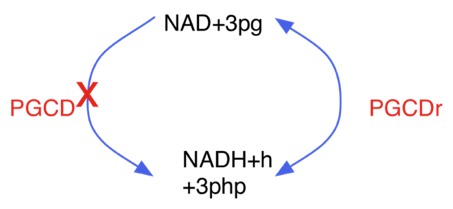
Turn off PGCD. Kept directionality of PGCDr as reversible.

Figure S1: Example of fixing cycles involving duplicate reactions.

**Case II: Lumped reactions**

In this case, multiple reactions in a pathway are lumped together to represent the overall conversion. If both the individual reactions and the lumped reaction are present in the model, they can potentially create thermodynamically infeasible cycles. The cycle can be broken by removing or turning off the lumped reaction and assigning proper annotation information to the individual reactions.

Example:

Aconitase (ACONT): cit_c[c] -> icit_c[c]

Aconitase (half-reaction A, Citrate hydro-lyase, ACONTa): cit_c[c] <=> h2o_c[c] + acon-C_c[c]

Aconitase (half-reaction B, Isocitrate hydro-lyase, ACONTb): icit_c[c] <=> h2o_c[c] + acon-C_c[c]

Solution:

The lumped reaction (ACONT) can be turned off.

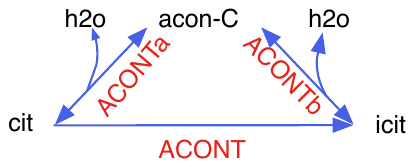

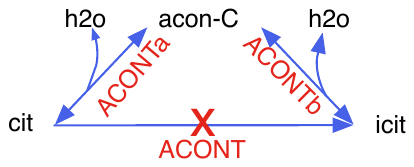

Figure S2: Example of fixing cycles involving lumped reactions.

**Case III: Cofactor specificity**

In this case, the same biochemical conversion is carried out by different cofactors in the model, while in reality the organism only uses one of the cofactors. If the cofactor specificity information is available, the reaction with non-specific cofactor can be removed or turned off.

Example:

D-Ribitol-5-phosphate NAD 2-oxidoreductase (DR1ORx ): nad_c[c] + dr5p[c] <=> h_c[c] + nadh_c[c] + ru5p-D_c[c]

and

D-Ribitol-5-phosphate NADP 2-oxidoreductase (DR1ORy): nadp_c[c] + dr5p[c] <=> h_c[c] + nadph_c[c] + ru5p-D_c[c]

both catalyzes the conversion of D-Ribitol-5-phosphate to Ribulose-5-phosphate in S. aureus.

Solution:

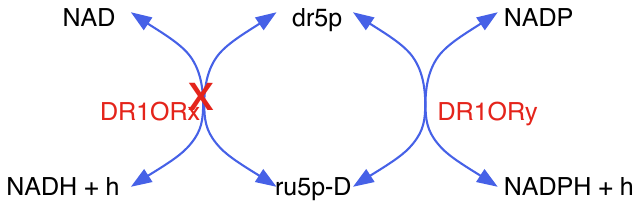

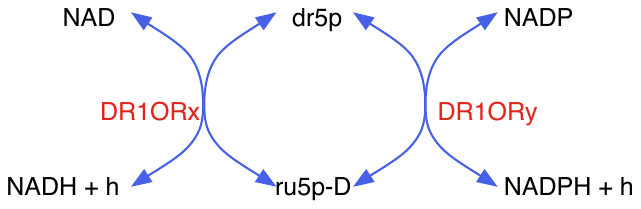
Upon extensive search for evidence in literature for cofactor specificity of S. aureus for this reaction, DR1ORx was turned off.

Figure S3: Example of fixing cycles involving non-specific cofactors.

2. **Consensus of *in vivo* essentiality information**

**Consensus on gene essentiality information**

Valentino et. al ^1^ hypothesized that large scale DNA sequencing combined with transposons can determine the essentiality and fitness criticality (non-essential but important for optimal growth) of genes. They determined gene essentiality by finding genes with fewer than 1% of the insertions expected from a random distribution and thereafter found 319 essential genes. The study quantified the effect of a mutation on fitness and found 106 genes that were important to fitness; these genes were defined as genes with 1-10% of the expected insertions. The study tested the mutants in a competitive growth environment with Brain Heart Infusion (BHI) Broth and found 420 essential genes. 30 of the 108 new essential genes were determined to be fitness compromised from before. The study found 91.1% of the previously reported essential genes, such as those by Chaudhuri et. al ^2^, to be essential. This suggests that this is an effective method for determining essential reactions. Of the remaining 8.9%, many were close to the essential cutoff, or were very far from being essential, suggesting that the results from the previous studies were inconsistent. One reason can be the presence of different false-positives within the different methods; and the 1% cutoff appears to have been arbitrarily determined, and genes important to fitness but not essential may fall within this cutoff. This work presumably incorporated true positives that were not included in Chaudhuri et. al ^2^ because Chaudhuri’s analysis made it hard to detect and classify essentiality in small genes or genes with few restriction sites. In general, Valentino was probably more accurate because they used 70,000 inserts compared to only about 350.

Santiago et. al ^3^ hypothesized that gene essentiality derived from transposon libraries can be affected by the high temperatures used to remove the plasmid delivery vehicle. As a result, previous methods found genes that were essential at high temperatures and/or at normal temperatures. They used a different methodology and was able to determine essentiality at 23°C, 30°C, 37°C, and 43°C. Santiago’s list of essential genes was determined at 30°C, instead of 37°C ^1^. Santiago had more inserts than Valentino (690,000 compared to 70,000), and also used a different methodology to determine essentiality, EL-ARTIST, instead of an arbitrary 1% cut-off. The increased number of inserts and EL-ARTIST allowed Santiago to determine if a gene was essential, had essential domains, or was non-essential. For the purpose of a metabolic model, the domain essential genes should be considered essential because knocking out the whole gene would kill the bacterium.

**Sources of Error**

There are two systematic sources of false positives in Chaudhuri et. al ^2^ and Valentino et. al ^1^. First, transposons can be incorrectly labeled essential if a polar effect from the transposon affects an essential gene immediately downstream. Second, the plasmid curing step requires high temperatures during a step of the experiment. This causes heat-essential genes to be incorrectly classified as essential genes. Santiago et. al ^3^ had evidence for likely false positive genes created by the polar effect. Santiago was able to upregulate and downregulate genes with transposons, enabling them to determine if a transposon was on an essential gene or just near an essential gene; these transposons were referred to as “erm” and “promoter”, respectively. However, the final data analysis uses a set of transposons referred to as “blunt” that affect only the downstream genes. 18 of the 20 essential genes found using the blunt methodology were found to be immediately upstream of an essential gene. Thus, genes found to be essential in the “blunt” dataset but non-essential in the “erm” and “promoter” sets are likely false positives, even if previous studies ^1,2^ found them to be essential. Santiago et. al ^3^ was unique because they were able to determine essentiality without a high temperature curing step. As a result, the study did not incorrectly label heat-essential genes as essential at 30°C. The study performed a test at 43°C to determine the heat-essential genes. They concluded that genes that were essential at 43°C but not 30°C are likely false positives. Valentino and Santiago randomly added transposons and then sequenced the junctions to determine which transposons remained after a number of generations.

The Nebraska Transposon Mutant Library (NTML) randomly generated transposon mutants and considered a gene to be knocked out if the transposon insertion was close to the 5’ end. NTML dataset was compared to the genes considered domain essential in Santiago et al ^3^. There appeared to be a relationship in which the domain essential genes have lower growth than normal. The average domain essential gene mutants have approximately 0.7 standard deviations of growth less than other mutants. The transposon may not disrupt the essential domain, allowing the gene to produce a slightly less functional protein. One would assume that there may be a relationship between genes considered fitness critical in Valentino et al ^1^ and the mutants with low growth in the mutant growth test. However, after looking at the fitness critical genes in agar conditions versus the growth in our mutant study, there appeared to be no correlation.

**Conclusions**

For metabolic modeling purpose, there are a few takeaways from the above discussion. First, the true list of essential genes may require a combination of many pools of knowledge. Genes found to be essential in any of the three data sets ^1-3^ should be considered essential unless 1) there was a growth mutant, 2) a gene was found to only be essential at 43°C, and 3) a gene was found essential with the blunt promoters but not under the other two methodologies. One exception is that domain essential genes were considered essential for the purposes of our model. This is because the mutant library may have only knocked out a portion of the gene, allowing it to still produce a functional protein. The second takeaway involves the number of fitness compromised mutants found in Valentino et al ^1^ and in the current study. These generally are not found in the *in silico* model; instead, the model generally shows full growth or no growth. This may suggest that some of the upper bounds in the model are too high, and the model may compensate for lost functionality by redirecting more flux through a pathway than possible *in vivo*.

3. **Growmatch Results**

*In silico* essential genes are found by turning off each gene individually and turning off the reaction(s) catalyzed by the gene by following the Boolean logic of the GPR relationships. *In vivo* essential genes were curated from multiple sources ^1-6^. Most of the essential genes were determined by randomly inserting transposons into Staphylococcus aureus and excluding the transposons which remained after growing the cells ^1-3^. An adaptation of data from two sources using antisense RNA was also used to determine essential enzymes and thus essential genes through the Boolean GPR relationships ^4-6^. The procedure for determining gene essentiality from the pool of literature is shown in Figure S4.

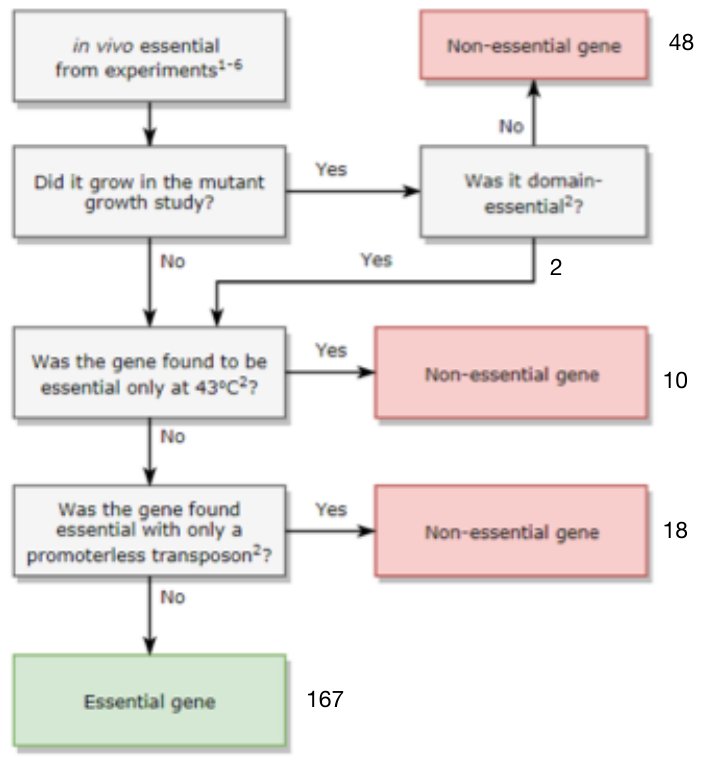

Figure S4. The methodology to determine gene-essentiality. The numbers around the boxes represent number of gene accepted/rejected from the essential gene list at different steps.

Genes found to be essential in any of the previous works were considered essential unless there was positive evidence suggesting the gene was non-essential ^1-6^. There were three types of positive evidence. First, mutants that were obtained from Nebraska’s Transposon Mutant Library ^7,8^ were not considered essential. Each individual mutant was grown in a 384-well plate to confirm that the mutant was able to grow. The least-fit mutant grew to an optical density (OD) of about 40% less than the average of the wild type control. As a result, all of these genes with a mutant were considered non-essential. An exception was made if the gene was found to be domain-essential ^3^. This is because the transposon may have inserted in a non-essential part of the gene, allowing a partially functional protein to be formed. Second, if the gene was found to be essential at only 43⁰C, then it is evident that the gene was incorrectly found to be essential in literature because of a high-temperature plasmid curing step in the processes used in the other literature sources ^3^. Third, if the gene was found to be essential using a promoterless transposon insert, but not with promoter-containing methodologies, then the gene is upstream of an essential gene, and other sources found it to be essential due to polar effects that disrupt expression ^3^.

After determining the list of *in vivo* essential genes, the growth and no-growth inconsistencies between experimental observations and the model predictions were reconciled and model performance was improved. Reactions and genes were categorized as G or NG, meaning that growth occurs when the gene or the corresponding reaction(s) was removed, or no growth occurs, respectively. GGs and NGNGs mean the model agrees with experimental evidence. GNGs are knockouts where the model predicts growth that does not occur in experiments, suggesting the model has spurious extra functionality; NGGs are knockouts upon which the model predicts no growth while experiments predict growth, suggesting the model lacks certain functionality. Growmatch is an optimization-based framework used to resolve metabolic models’ growth predictions with experimental evidence^9^. Growmatch consists of two algorithms: GrowmatchNGG and GrowMatchGNG.

**GrowmatchNGG:**

GrowmatchNGG was used to resolve NGGs. The algorithm turns off an NGG gene, and then maximizes growth by adding the minimal amount of reactions (minimum one to maximum four was allowed in this work) from a database of reactions. These solutions come from three sources: 1) The backwards directions of irreversible reactions (because the reversibility of the reaction may be uncertain) 2) Transport reactions of metabolites ( either diffusion through the cell membrane or via a non-specific transporter), and 3) The reactions from taxonomically similar organisms (*E. coli* and *B. subtilis*) from the BIGG database ^10^. Growmatch suggested various solutions (a set of one to four reactions) to solve most of the NGGs. Solutions were added one by one and checked to ensure they do not invalidate any NGNGs or create any new thermodynamically infeasible reactions cycles. Solutions were prioritized if they resolved NGGs in the central metabolism or amino acid biosynthesis. The rest of the solutions were arranged in order of increasing sum of ranks. The reactions were ranked from one (the most likely) to four (the least likely). Database reactions from *Staphylococcus aureus* models and the backwards direction of irreversible reactions were given a ranking of one. Reactions from the phylum Firmicutes were given a ranking of two 2. Reactions from other bacteria were given a ranking of three 3. Reactions from a different domain were given a ranking of four. Only one transport reaction was suggested, and it was given a ranking of 3 because it has been found in *E. coli^2^*.

Table 1: Ranking scheme for GrowMatchNGG solutions.

| Rank | Origin | Rationale |
| --- | --- | --- |
| 1 | *S. aureus* or changing direction | The values of ΔG have a low certainty in the cell, so the reactions could be reversible. |
| 2 | Phylum Firmicutes | Same lower taxonomic group |
| 3 | Other Bacteria | Same higher taxonomic group |
| 4 | Other Kingdoms | These reactions are known but do not have any evidence of existing in bacteria. |

After checking to ensure the solutions did not invalidate NGNGs or create any new thermodynamically infeasible cycles, one solution for each NGG was added to the model.

**GrowmatchGNG :**

GrowmatchGNG was used to suggest reactions or reaction directions to remove to resolve GNGs. GrowmatchGNG turns off a GNG gene and attempts to minimize the maximum growth by turning off one or multiple reactions (the candidate solutions). GrowMatchGNG produces multiple solutions that can resolve the GNG. In order to minimize the solution space for two reaction knockouts, reactions from iSB619 were not considered for removal. This was justified because the reactions in iSB619 had a gene associated with the reaction or a rationale for each reaction ^11^. The solutions were pruned by ensuring they did not violate any GGs.
